# Compartmentalized Cytoplasmic Flows Direct Protein Transport to the Cell’s Leading Edge

**DOI:** 10.1101/2024.05.12.593794

**Authors:** Catherine G. Galbraith, Brian P. English, Ulrike Boehm, James A. Galbraith

## Abstract

Inside the cell, proteins essential for signaling, morphogenesis, and migration navigate complex pathways, typically via vesicular trafficking or microtubule-driven mechanisms ^1–3^. However, the process by which soluble cytoskeletal monomers maneuver through the cytoplasm’s ever-changing environment to reach their destinations without using these pathways remains unknown. ^4–6^ Here, we show that actin cytoskeletal treadmilling leads to the formation of a semi-permeable actin-myosin barrier, creating a specialized compartment separated from the rest of the cell body that directs proteins toward the cell edge by advection, diffusion facilitated by fluid flow. Contraction at this barrier generates a molecularly non-specific fluid flow that transports actin, actin-binding proteins, adhesion proteins, and even inert proteins forward. The local curvature of the barrier specifically targets these proteins toward protruding edges of the leading edge, sites of new filament growth, effectively coordinating protein distribution with cellular dynamics. Outside this compartment, diffusion remains the primary mode of protein transport, contrasting sharply with the directed advection within. This discovery reveals a novel protein transport mechanism that redefines the front of the cell as a pseudo-organelle, actively orchestrating protein mobilization for cellular front activities such as protrusion and adhesion. By elucidating a new model of protein dynamics at the cellular front, this work contributes a critical piece to the puzzle of how cells adapt their internal structures for targeted and rapid response to extracellular cues. The findings challenge the current understanding of intracellular transport, suggesting that cells possess highly specialized and previously unrecognized organizational strategies for managing protein distribution efficiently, providing a new framework for understanding the cellular architecture’s role in rapid response and adaptation to environmental changes.

## Main

In the dynamic and crowded cellular environment, how proteins navigate the cytoplasm to reach their destinations without traveling in vesicles or hitching a ride on motors remains largely enigmatic. We have uncovered a critical role of the actin-myosin network in forming a semi-permeable barrier that not only segregates but also actively targets both actin and other proteins through a specialized compartment. This compartment harnesses contraction-induced, molecularly non-specific fluid flows to propel proteins precisely to local areas where new filament growth occurs at the cell’s protruding edges. Our findings reveal a sophisticated cellular strategy for managing protein transport. They provide a novel perspective on the cell’s ability to reconfigure its internal architecture for precise and efficient protein localization during essential morphological changes and migration.

To elucidate the dynamics of protein transport across the cytoplasm, our initial experiments focused on tracking how depolymerized actin monomers move toward the cell front for repolymerization. The process of actin polymerization, conserved for a billion years ^7^, is biochemically well-understood and can even be reconstituted in vitro ^8^. Despite this, the mechanisms by which actin monomers navigate through the cytoplasm to contribute to cell remodeling and migration remain elusive. Mass balance necessitates that depolymerized monomers must be repolymerized; however, visualizing actin’s fast, directional transport against the background of treadmilling actin networks is inherently difficult. Although indirect measures have suggested that actin moves via fluid flow ^9, 10^, this mechanism is so poorly understood that it is often represented in models by a simple forward-directed arrow ^11^. Here, we use direct measurements to identify a complex interplay of biochemical and mechanical forces that manages protein distribution for rapid shape changes.

To measure the velocity of actin movements, we utilized photobleaching to mark the treadmilling network of NG108 cells expressing EGFP-actin, specifically targeting the lamella’s rear to maximize the monomers’ travel distance to the leading edge, where depolymerization predominantly occurs ^12^. Shortly after bleaching, a distinct thin black line became visible at the cell front within a few seconds (Fig 1a), indicating the rapid anterograde movement of actin monomers from the bleached zone. The pronounced thinness of this line is likely due to the higher concentration of barbed filament ends, which are available for polymerization ^13, 14^, and the integration of some of the depolymerized monomer into other parts of the dendritic network as it moves forward^15^. By measuring the distance between the initial bleach and the appearance of this black line and dividing by the elapsed time, we calculated the velocity of anterograde transport to be 3.9 ±1.7 μm/s, nearly fifty times faster than retrograde flow, measured by the rearward movement of the original bleach line (Fig 1d, Extended Data Fig 1).

**Figure 1:**
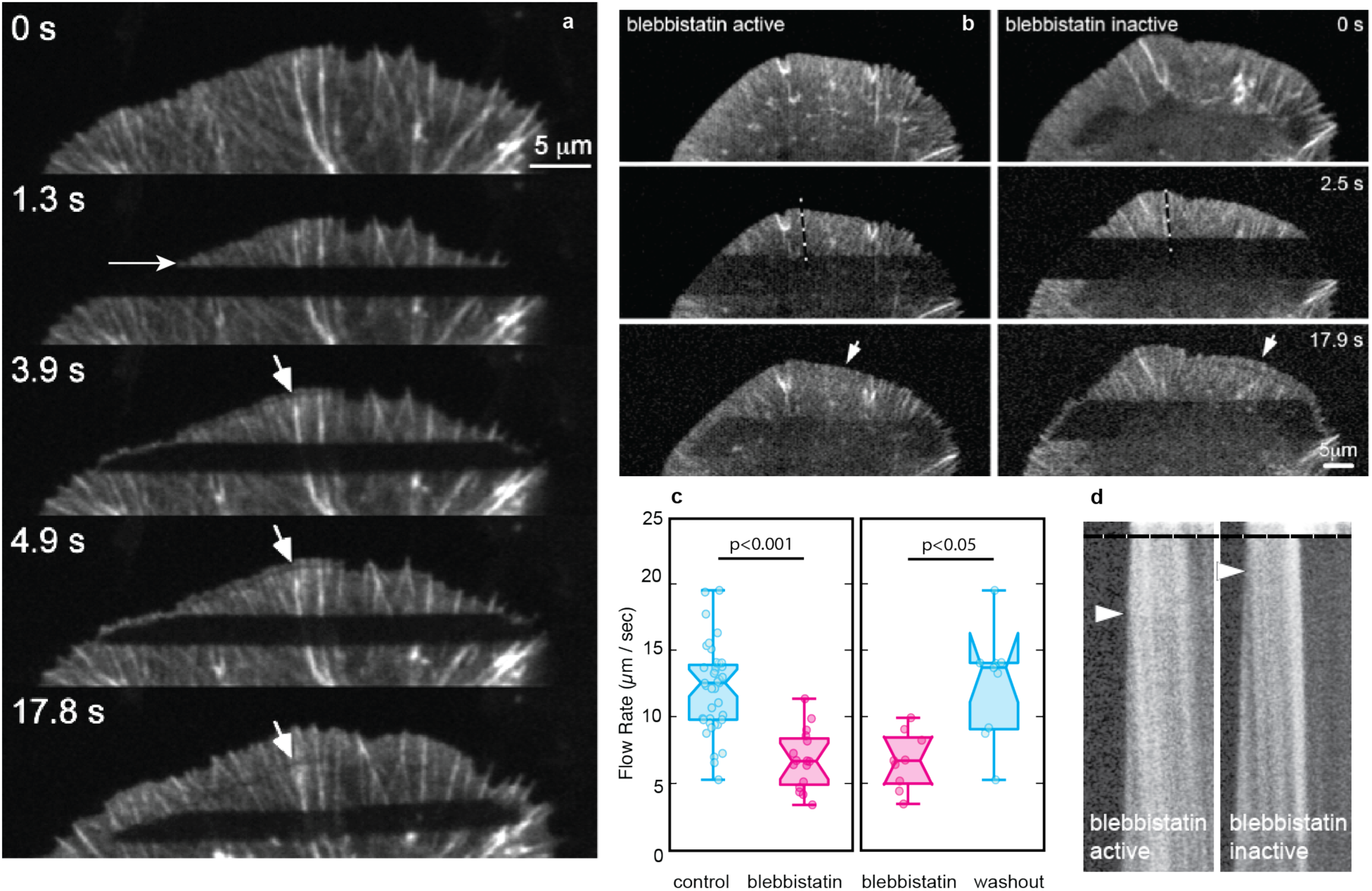
Transport of depolymerized actin to the polymerizing front is facilitated by myosin contraction. **a,** The EGFP-actin network of NG108 cells was rapidly bleached between 0.3 and 1.3s. At 3.9s, bleached actin monomer from the network has been transported (recycled) to the front of the cell, repolymerized at the leading edge, and traveled rearward (thin dark line indicated by arrow). **b**, Forward actin transport is delayed when myosin II ATPase is inhibited with blebbistatin in cells expressing Apple-actin. Photonic washout - inactivation of blebbistatin with 488 nm light - rescues the speed of rapid actin forward transport (indicated by the return of the dark line at the cell front (arrow). **c**, Forward (anterograde) flow rate in control and blebbistatin treated cells and treated matched photonic washout experiments (n=38 control, 16 blebbistatin (20 μM), 9 photonic washout, p values 2 tailed comparisons). **d,** Kymographs of the lines indicated in (**b**) indicate the time until actin from the wide bleach band appears as the narrow band at the cell front (arrow). Scale bars, 5 μm. All panels represent at least three independently performed experiments.

To elucidate the mechanisms behind this fast anterograde transport, we focused on myosin II, which has been shown not to affect transport ^16^ but has also been suggested to generate a pressure gradient through contraction at the cell rear or around the cell body that facilitates transport^9, 17^. To address these conflicting results, we inhibited myosin II activity by treating cells expressing Apple-actin with a wavelength-dependent version of the myosin II ATPase inhibitor, blebbistatin ^18^. Post-treatment, we observed a diffuse line that appeared near the cell edge more slowly than in control cells, illustrating reduced transport efficiency (Fig1b, LHS). Subsequent photonic inactivation of blebbistatin with 488 nm light within the same cell restored the line’s sharpness and the speed of anterograde movement (Fig1b, RHS), suggesting that active myosin contraction is crucial for rapid protein transport. Despite the retrograde speed of the network remaining constant, the forward speed was reduced by 2-4X fold under myosin inhibition by either blebbistatin or the Rho-kinase inhibitor Y-27632 ^19^ (Extended Data Fig 1b, c). We modeled the data according to a simple 1D diffusion equation, confirming that inactive myosin leads to transport speeds comparable to diffusion alone ^20^. Calculation of the Péclet number, Pe, yielded a value of 3.5; values >1 indicate that advection dominates diffusion ^1, 21^ (Methods).

While earlier modeling efforts proposed such an advective forward flow to transport protein forward, these studies could not experimentally visualize a flow ^9, 10^. To bridge this gap, we borrowed methodologies from fluid mechanics, specifically adapting the technique for measuring the dissipation of material from a continuous point source ^22^. In this experiment, we utilized CAD cells, which feature a broad lamella less dominated by filopodia compared to NG108 cells and expressed photoactivatable green fluorescent protein (PaGFP)-tagged actin, which remains dark until illuminated by UV light ^23^. We then introduced a focused UV beam to the field of view and simultaneously imaged the activation to track the dispersion of actin. This method, akin to an inverse Fluorescence Loss in Photobleaching (FLIP) experiment ^24^, which we term ‘FLOP’ for Fluorescence Leaving the Original Point, revealed distinct transport dynamics. When activating a spot within the cell body, fluorescence dispersed symmetrically, consistent with diffusion. In contrast, activation within the lamellipodial region produced an asymmetric plume of fluorescence stretching toward the leading edge, mirroring the expected pattern of a fluid flow’s analytical solution (Fig 2a, b, Extended Data Fig 3) ^22^. This plume continued until the fluorescently labeled actin monomers reached and were incorporated into the advancing actin network at the cell front (Fig 2a, d).

**Figure 2:**
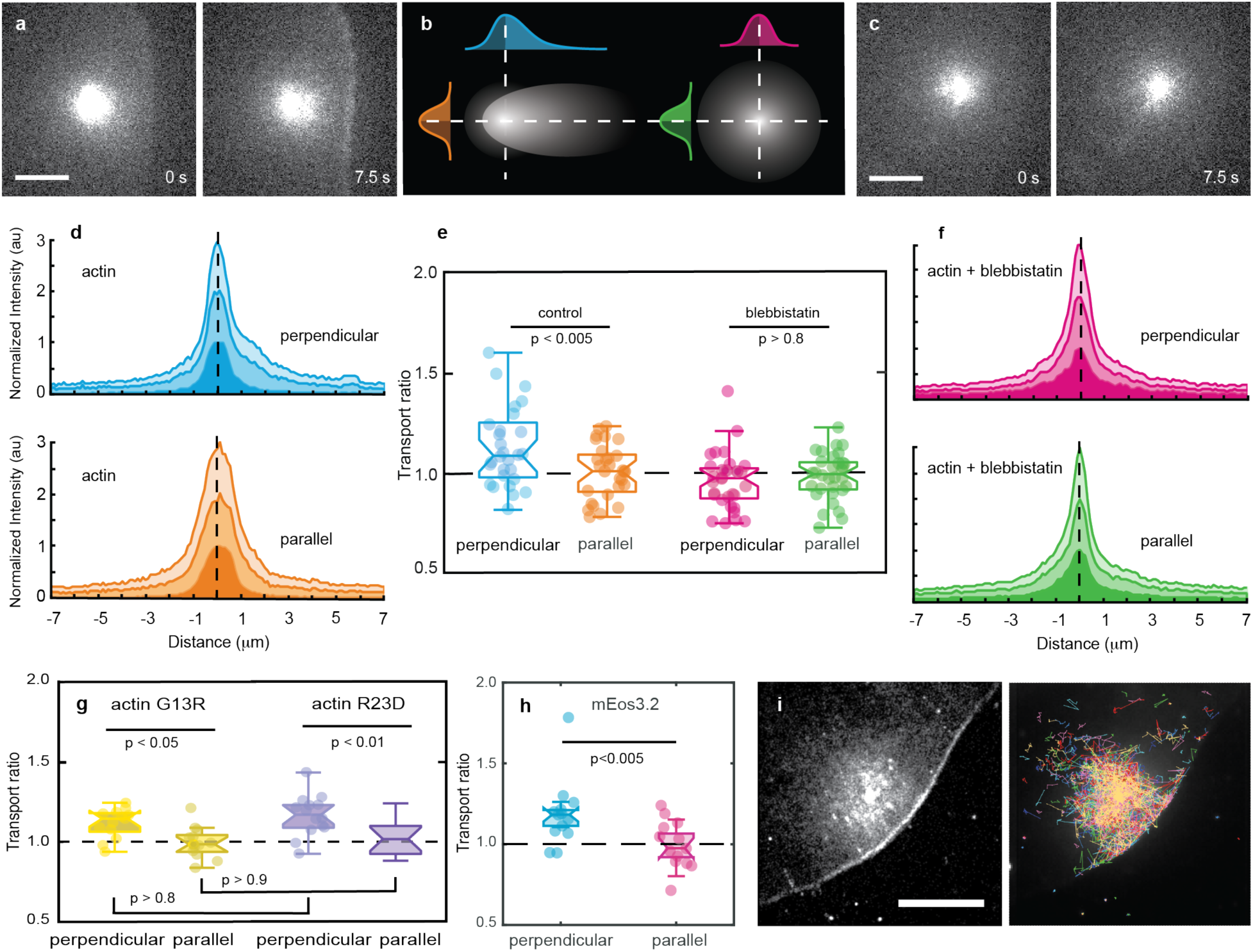
Myosin mediates the forward flow of actin. **a**, Photo-activation of a diffraction-limited spot in a CAD cell expressing PaGFPactin shows asymmetric movement toward the front of the cell. **b**, Graphic illustrating how asymmetric spread is analyzed from fluorescence intensity perpendicular and parallel to the cell leading edge. The transport ratio is defined as the area under each half of the intensity curve. **c,** Photo-activation of a diffraction-limited spot in a CAD cell expressing PaGFPactin shows asymmetric movement toward the front of the cell. **d,** Line-intensity plots indicate the asymmetry in the control cell in (**a**) is only in the direction of the cell leading edge. t=1.5, 3, 7.5 s after the initiation of spot activation. **e**, Box and whisker plots, including all individual data points, indicate fluorescence dispersion toward the leading edge (perpendicular) in control cells. (n= 29 control, 31 blebbistatin (0.2 μM). **f,** Line-intensity plots indicate the symmetry in blebbistatin treated cell in (**c**). t=1.5, 3, 7.5 s after the initiation of spot activation. **g**, Asymmetry analysis as described in b for the spread of actin mutants that cannot be incorporated into the network (n=13, G13R, 14, R62D). **h**, Asymmetry analysis for free dye, mEos3.2 (n=13). **i**, Sum of x single-molecule frames and single molecule tracking from spot activation of mEos3.2-R62D. p values two-tailed t-tests in (**e**) and (**h**), Scheffe post-Anova test. Scale bars, 5 μm. All panels represent at least three independently performed experiments.

This experimental setup facilitates quantification by measuring fluorescence intensity distribution both forward and rearward from the central line of the activation point orthogonal to the cell edge (Fig 2b). Calculations parallel to the cell edge served as a control for symmetric dispersion, indicative of diffusion (Fig 2b). Inhibition experiments using the wavelength-independent version of the myosin II ATPase inhibitor, nitro-blebbistatin, and the Rho-associated protein kinase (ROCK) inhibitor, Y27632, consistently produced symmetric fluorescence profiles in both directions (Fig 2c, e, f, Extended Data Fig 1d, e). These data indicate that in the absence of active myosin II, the transport mechanism shifts towards diffusion, further substantiating the role of myosin II-dependent advection in driving the polymerizable monomer toward the leading edge.

To test whether the observed asymmetry resulted from monomer incorporation into the network at the cell front, we compared the dispersion patterns of two actin mutants, G13R and R62D, which cannot be incorporated into the network. Surprisingly, both mutants showed asymmetric distributions similar to wild-type actin. Notably, while G13R can bind profilin— involved in actin assembly and organization at the cell edge—R62D binds to cofilin, another regulatory protein ^25^. The similarity in the asymmetry observed with these mutants and actin suggests that neither direct incorporation into the network at the leading edge nor interaction with profilin is essential for generating the observed transport asymmetry (Fig 2g) ^26^.

These results led us to explore whether the observed transport mechanism extends beyond actin recycling to serve as a generic, molecularly non-specific system for relocating various proteins to the polarized leading edge of cells. Initial attempts to track the distribution of untagged PaGFP were hindered by its inadequate ON/OFF contrast ratio ^27^, prompting us to switch to the higher-contrast photoactivatable probe mEos3.2, known for its utility in single-molecule microscopy and tracking ^28^. Subsequent experiments demonstrated that both mEos3.2 and mEos3.2-R62D were propelled toward the leading edge (Fig 2h, i), exhibiting patterns of accumulation reminiscent of those observed with small molecules in migrating keratocytes, attributed to pressure-driven flow ^9^. To quantify the movement, we conducted single particle tracking of mEos3.2-R63D and JF549 Halo-G13R. We calculated momentum scaling spectrum analysis (MSS) for individual trajectories, which revealed that a significant percentage of the molecular population moves exhibited facilitated diffusion (Extended Data Fig 4) ^29^. This observation aligns with simulations that required a subset of molecules to undergo high-speed flow to replicate the observed rates of protein transport across different cellular regions ^10^. Our data suggest that this transport mechanism can facilitate the forward movement and precise localization of various proteins, supporting the existence of a broadly applicable, advection-driven protein transport system at the cell front.

To determine if advection was also the dominant transport mode within the cell body, we extended our investigations of PaGFP-actin dispersion in longer time-course FLOP experiments. These experiments revealed that fluorescence rapidly diffused throughout the cell body, indicating a distinct transport mechanism compared to the lamella (Fig 3a). Notably, when we activated the network in the lamella region, the fluorescent actin primarily remained localized, with minimal spillover into the cell body (Fig 3a). A critical difference was observed when the activation spot was positioned at the actin-myosin boundary; here, fluorescence spread in both directions, highlighting the boundary’s role as a semi-permeable barrier (Fig 3b, Extended Data Fig 5a). This barrier effectively segregates the lamella, where advection dominates, from the rest of the cytoplasm, where diffusion appears to be the primary mode of transport. These findings suggest the existence of a specialized compartment at the cell front, distinct in its transport dynamics from the cell body.

**Figure 3.**
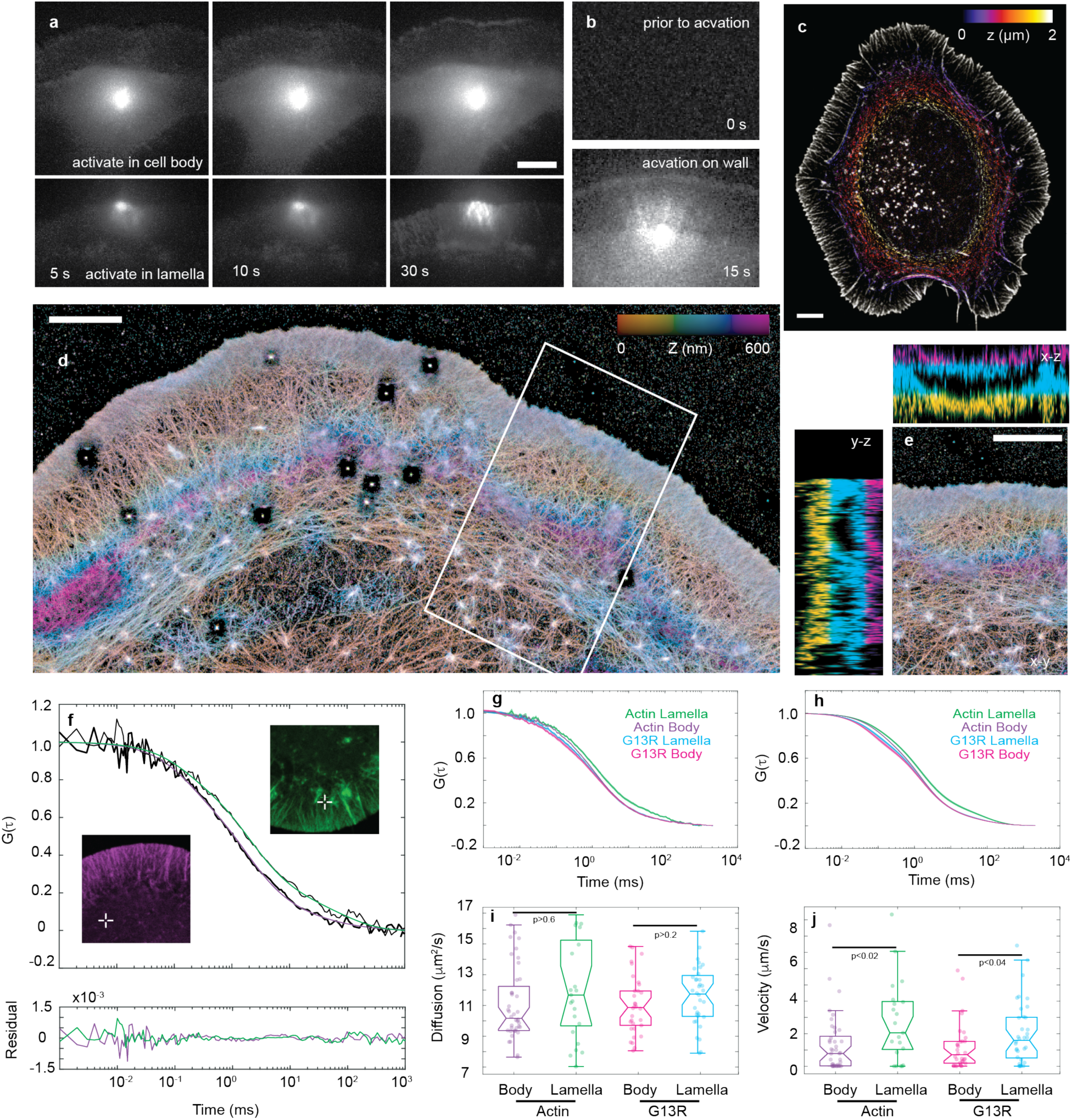
Actin cytoskeleton creates an isolated compartment with a separate transport system at the cell front. **a,** Spot activation of PaGFP actin in a CAD cell demonstrating rapid filling of fluorescent material throughout the cytoplasm. At 30 seconds, actin has traveled into the lamella compartment and is incorporated into the leading edge; however, the time to incorporate is much slower than a linear scaling of the data from Fig 2 due to increased distance from the edge. **b,** When the spot is placed on the lamella-cell body barrier, actin rapidly spreads in both directions. **c,** Maximum projection of a 3D SIM image of mEmerald MLC, Alexa 647 Phalloidin labeled CAD cells, color-coded for height, indicates that actin-myosin is arranged vertically between the lamella and the cell body. **d,** iPALM image of f-actin rendered at 15 nm isotropic resolution, color-coded for height. Blue and magenta colors in the middle of the inset box indicate the increase in actin arcs at the rear of the lamella. **e,** y-z, and x-z inset panels from (**d**) indicate a compartment between the leading edge and the actin arcs at the base of the lamella. The color code is the same as in (**d**). **f**, Representative FCS autocorrelation curves (black lines) of Halo Actin labeled with JF549 taken from the lamella and cell body. The fit to a model of two diffusion coefficients and flow is shown in color, with the accuracy of fit indicated by the residual panel**. g,** Data mean ± SE from Halo-actin or Halo-G13R in the lamella or the cell body (n =21 actin lamella, 40 actin cell body, 32 G13R lamella, 32, G13R cell body). The shift of the actin lamella curve to the right is indicative of flow, but the other three curves overlap. **h,** Mean ±SE of model fit to data in (**g**). **i,** Primary diffusion is not significantly different when actin can or cannot bind the network and when measured in the cell body or the lamella. **j**, The flow velocity is very small in the cell body and statistically different in the lamella. p values two-tailed t-tests t. Scale bars, 5 μm. All panels represent at least three independently performed experiments, except iPALM, which was repeated once.

To define the structure of the barrier, we employed 3D structured illumination microscopy (3D-SIM). We discovered a dense vertical concentration of actin and myosin at the lamella-body boundary in both CAD and NG108 cells (Fig 3c, Extended Data Fig 5b, c). Building on this, we utilized interferometric photoactivated localization microscopy (iPALM), which allowed us to reconstruct the actin nanoarchitecture with a 15 nm isotropic resolution. iPALM revealed a vertical wall that effectively segregated the lamella from the cell body (Fig 3d). This structural configuration confirms the existence of a distinct physical compartment that isolates the lamella’s active transport activities from the more passive transport within the cell body (Fig 3e, Extended Data Fig 6).

The integration of spot activation data and the discovery of a semi-permeable barrier between the lamella and the cell body underscored the presence of distinct transport mechanisms in each compartment. We then sought to quantify these dynamics with fluorescent correlation spectroscopy (FCS), which tracks fluctuations in fluorescence intensity over time and computes their autocorrelation function to reveal underlying molecular movements ^30^. Our FCS analysis in CAD cells expressing either Halo-tagged actin or the G13R mutant labeled with JF549 yielded autocorrelation functions that fit well with a two-component diffusion model with flow, indicating two distinct molecular behaviors: free monomer diffusion and monomer interaction with other proteins, influenced by fluid flow, as indicated by the magnitude of the fit residues (Fig 3f) ^30–32^. The right shift of the actin curve compared to the cell body suggested the presence of fluid flow in the lamella. The tight clustering mean curves and their fits make it difficult to distinguish between the different conditions (Fig 3g, h, Extended Data Fig 7). However, the flow and diffusion dynamics analysis separated the data. We found the diffusion rates were similar across both compartments, aligning with established literature values (Fig 3i), but the fluid flow velocities were notably higher in the lamella, approximately double that observed in the cell body for both actin and the G13R mutant (Fig 3j). This finding decisively points to forward-directed advection as the predominant transport mechanism in the lamella, contrasting with the diffusion-dominated transport observed in the cell body.

These findings help reconcile previously conflicting models that proposed the existence of advective flow. While a minor amount of fluid does flow within the cytoplasm, monomer movement in the cell body is predominantly driven by diffusion. However, the transition of monomers between the cell body and the lamella is significantly slowed by the semi-permeable actin-myosin barrier. This compartmentalized advection model underscores its critical role in establishing the specialization of the leading edge of migrating cells, where actin polymerization catalyzes protrusion, alters adhesion receptor conformations, and forms connections to the extracellular matrix that anchor and propel the cell forward. To validate this model further, we analyzed the movement of the Arp3 component of the Arp2/3 complex and paxillin (Fig 4a), which are essential for edge extension and adhesion formation. Using a point source to activate the protein continuously, we found that they also exhibited adventive transport toward the leading edge. This indicates that crucial proteins required for cellular migration and adhesion also utilize this accelerated transport mechanism, moving faster than the typical speed of diffusion, to get where they need to be.

**Figure 4.**
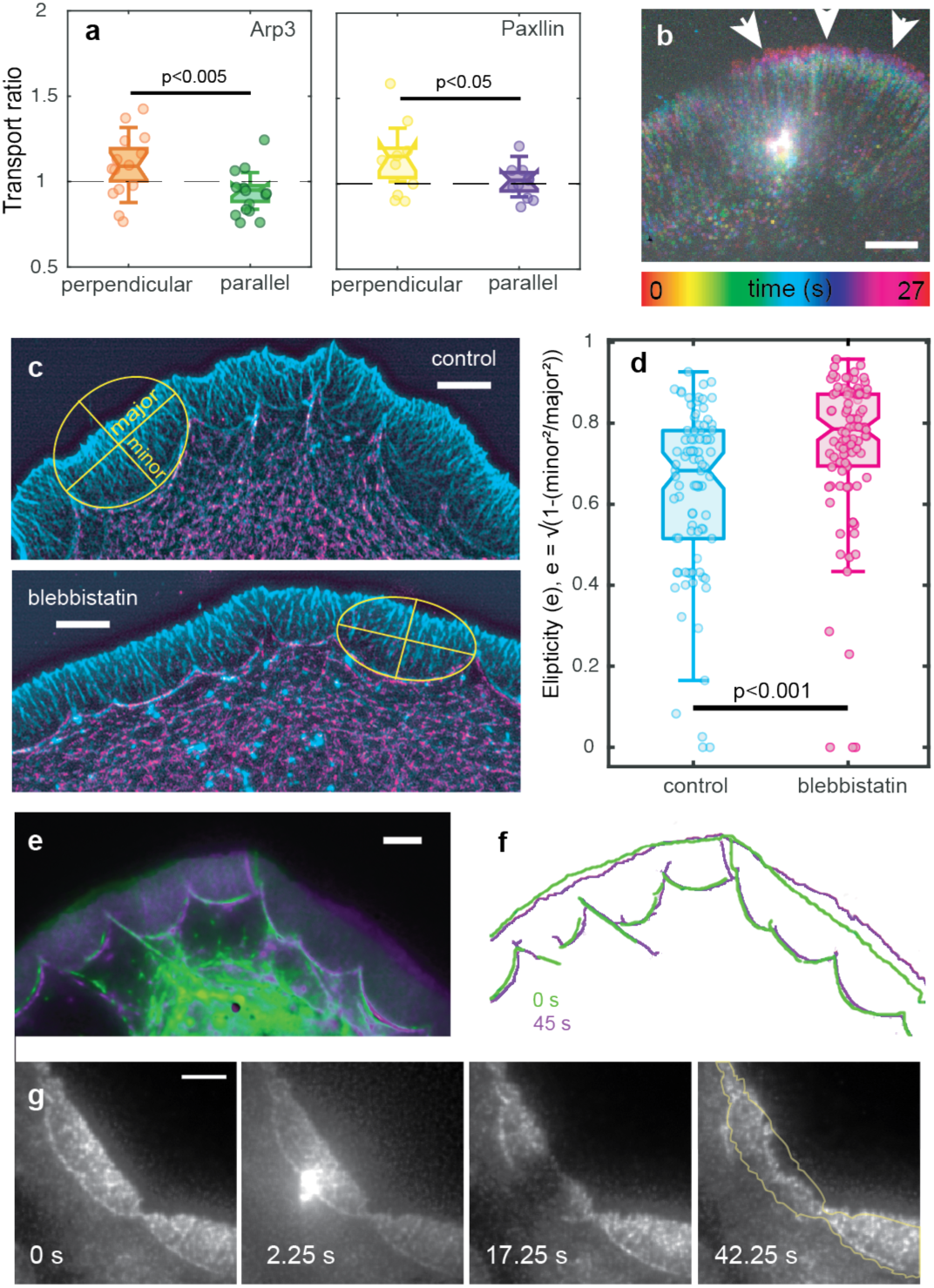
The myosin wall targets polymerizable proteins to advancing regions of the leading edge. **a,** Asymmetry analysis of spot activation of mEos3.2 Arp3 and paxillin in CAD cells illustrate that both are targeted toward the leading edge, where polymerization and adhesions initiate (n=9 Arp3, 9 paxillin). **b,** Temporal color-coded spot activation of mEos3.2 actin in an NG108 cell advancing preferentially on the right, where activated actin is targeted. **c,** Maximum projection of 3D SIM images of Alexa 647-phalloidin and mEmerald MLC in control cells and cells treated with 0.2 μM nitro-blebbistatin before fixation. The actin-myosin arcs at the border between the lamella and the cell body were fit to ellipses. The ellipticity was calculated as shown and quantifies the broader, flatter arcs in the blebbistatin-treated cells (n=23 control cells, 87 measurements, 19 blebbistatin cells, 100 measurements). **d,** Overlay of two different time points of mNeon MLC image (t=0 green, and t=45 s purple illustrating that the edge retracts on the left (green at the edge) and extends on the right (purple on the edge). **e,** Contour of same two-time points in (**d**), illustrating that on the side that extends, the curvature of the arcs increases and tilts toward the local expansion while the arcs flatten in the side that is retracting. **f,** CAD cell expressing mEmerald MLC with a high-power localized beam of 405 light positioned over one arc, causing the arc to be disrupted and only the leading edge in front of that single arc to collapse. p values two-tailed t-tests t. Scale bars, 5 μm.

We then conducted additional spot activation experiments to explore the specificity of the transport mechanism in relation to the irregular advancement of different regions of the leading edge. Using NG108 cells, we spot-activated mEos3.2-actin and observed that, instead of dispersing uniformly toward the front of the cell, actin preferentially moved towards and incorporated into actively advancing regions, suggesting a directional bias in the flow (Fig 4b). This pattern was corroborated in 3T3 fibroblasts using locally activated PaGFP-actin, confirming that the advection mechanism targets advancing cell regions (Extended Data Fig 8). Further investigations in CAD cells revealed that the actin-myosin wall at the rear of the lamella, which forms well-defined arcs, plays a crucial role in this targeting. The arcs demonstrated dynamic changes in curvature that correlated with the directional advancement or retraction of the edge (Fig 4e). Notably, the curvature, quantified as ellipticity, flattened when myosin activity was inhibited with blebbistatin, coinciding with a decrease in fluid flow (Fig 4c, d). Additionally, disrupting a single arc by laser ablation led to the local collapse of the leading edge in front of the arc (Fig 4f). These findings indicate that the actin-myosin arcs direct depolymerized monomers from the treadmilling network to specific locations along the edge and regulate the speed and direction of transport.

## Discussion

As the cytoskeleton remodels and alters its shape, proteins travel across the cytoplasm to get where they are needed. Here, we provide direct observation of cytoskeletal remodeling creating fluid flow, driven by myosin contraction, to target proteins to sites of polymerization and scaffold assembly. While traditional models suggest that myosin contraction around the cell body facilitates diffusion by generating fluid flow, our findings using fluorescence flow, FCS, iPALM, and single particle tracking paint a different picture. Myosin contraction indeed generates fluid flow, but this flow is not merely a simple diffusion facilitator; instead, it actively directs transport within a specific cellular compartment, targeting locally advancing regions of the leading edge.

We also found that, in conjunction with myosin, the actin cytoskeleton creates a barrier separating the lamella from the rest of the cell body, creating what functions as a novel pseudo-organelle, bordered by cellular membranes on all sides except at the actin-myosin wall. The compartmentalization sheds new light on earlier studies, suggesting two sources of actin monomer ^33^. Our data indicate that they should be reinterpreted as crossing the semi-permeable actin-myosin barrier, which ensures that transport within each compartment is optimized for specific cellular functions.

Having a separate compartment at the front allows for much faster and finer control of protein transport as migrating cells protrude, probe their environment for ECM, and form adhesions to serve as anchors to move themselves forward. Protein only needs to move short distances for rapid redeployment to participate in protrusion and adhesion formation. This localized, rapid movement directs and concentrates proteins at protruding regions of the leading edge, conducive to specific biochemical reactions, such as actin polymerization, and influences integrin conformation at the protruding edge ^34^. Additionally, the dynamic curvature of the actin-myosin wall under myosin contraction finely tunes the direction and rate of protein delivery along the leading edge. Thus, the architecture of the actin cytoskeleton enables the process of treadmilling to facilitate the supply of its own monomer for depolymerization during a steady state and rapid shape change and migration.

In summary, our findings reconceptualize the front of the cell as a dynamic, ‘membrane-less’ organelle ^35^, which perpetually directs polymerizable proteins to precisely where they are most needed. By strategically targeting advancing areas along the leading edge, the myosin arcs modulate the polymerization-dependent conformational activation of integrin, priming it for effective ECM binding. This targeted protein delivery enhances the cell’s ability to rapidly adapt and respond to extracellular cues, underscoring the sophisticated orchestration of cellular structures and functions that drive cellular motility and interaction with the environment.

## Acknowledgments

All authors contributed to drafting the manuscript and gratefully acknowledge Luke D. Lavis for his gifts of JF dyes and sponsorship of the Janelia Visitor Program (CGG). This work was supported by NIH grant GM117188 (CGG), NSF award 171636 (JAG), and the Howard Hughes Medical Institute (BPE and UB). iPALM work was partly supported by a Janelia Advanced Imaging Center award. SIM imaging was partly supported by a Core Research Facilities Grant from OHSU School of Medicine.

---

**Extended Data Fig 1.**
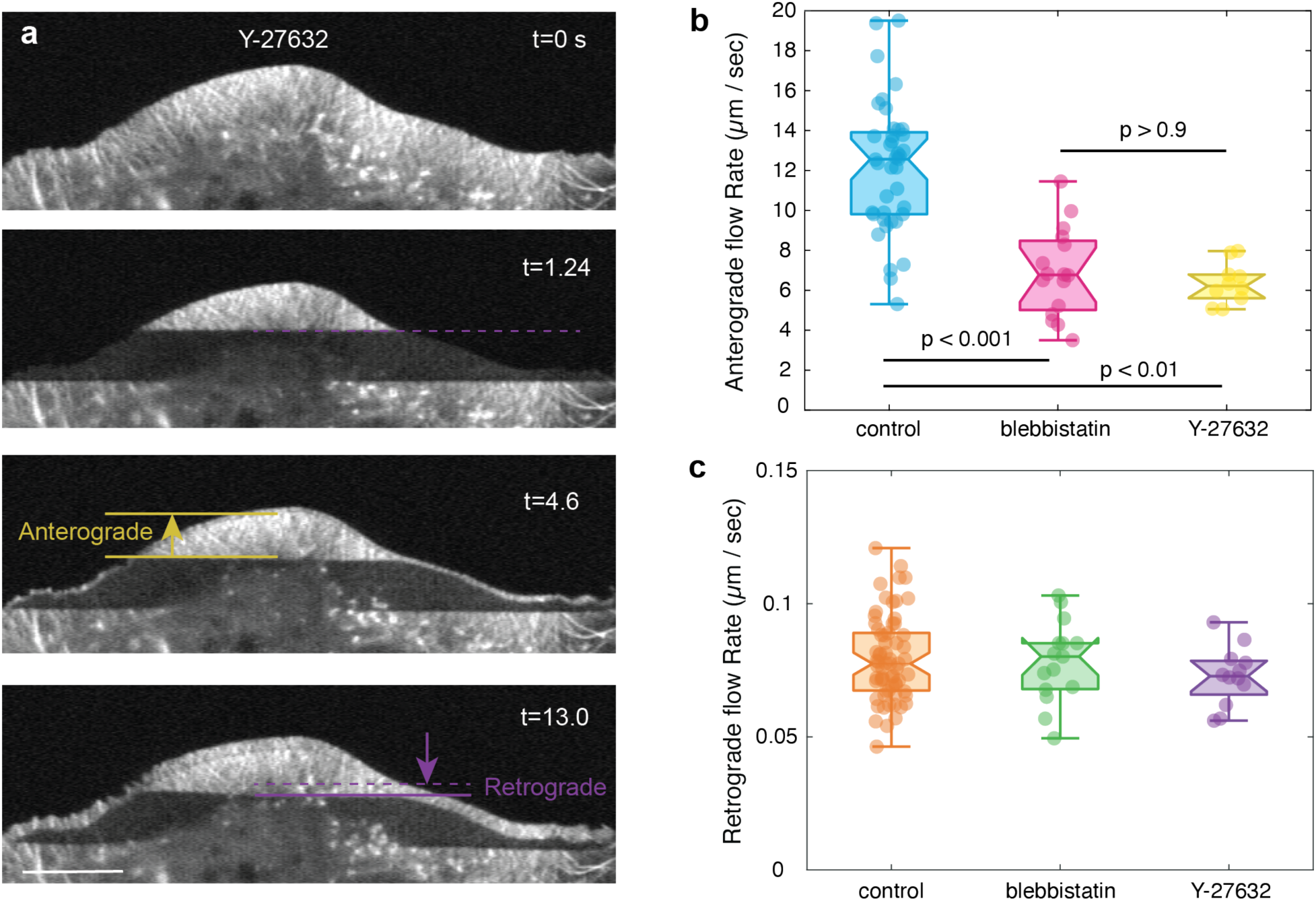
Myosin inhibition does not affect treadmilling or the network retrograde flow rate. **a**, A Typical NG108 cell expressing EGFP-actin rapidly bleached while under Rho Kinase inhibitor Y-27632 (10 μM), showing a delayed appearance of a thin line at the edge. Time series annotated to indicate the methodology for measuring flow. **b**, Anterograde flow rate of Y-27632 treated cells is similar to blebbistatin treated cells (n=10 Y-27632, control and blebbistatin anterograde flow from Fig1**c**. (p values Scheffe post ANOVA test). **c**. None of the treatments affected the retrograde flow rate. Scale bars, 5 μm. All panels represent at least three independently performed experiments.

**Extended Data Fig 2.**
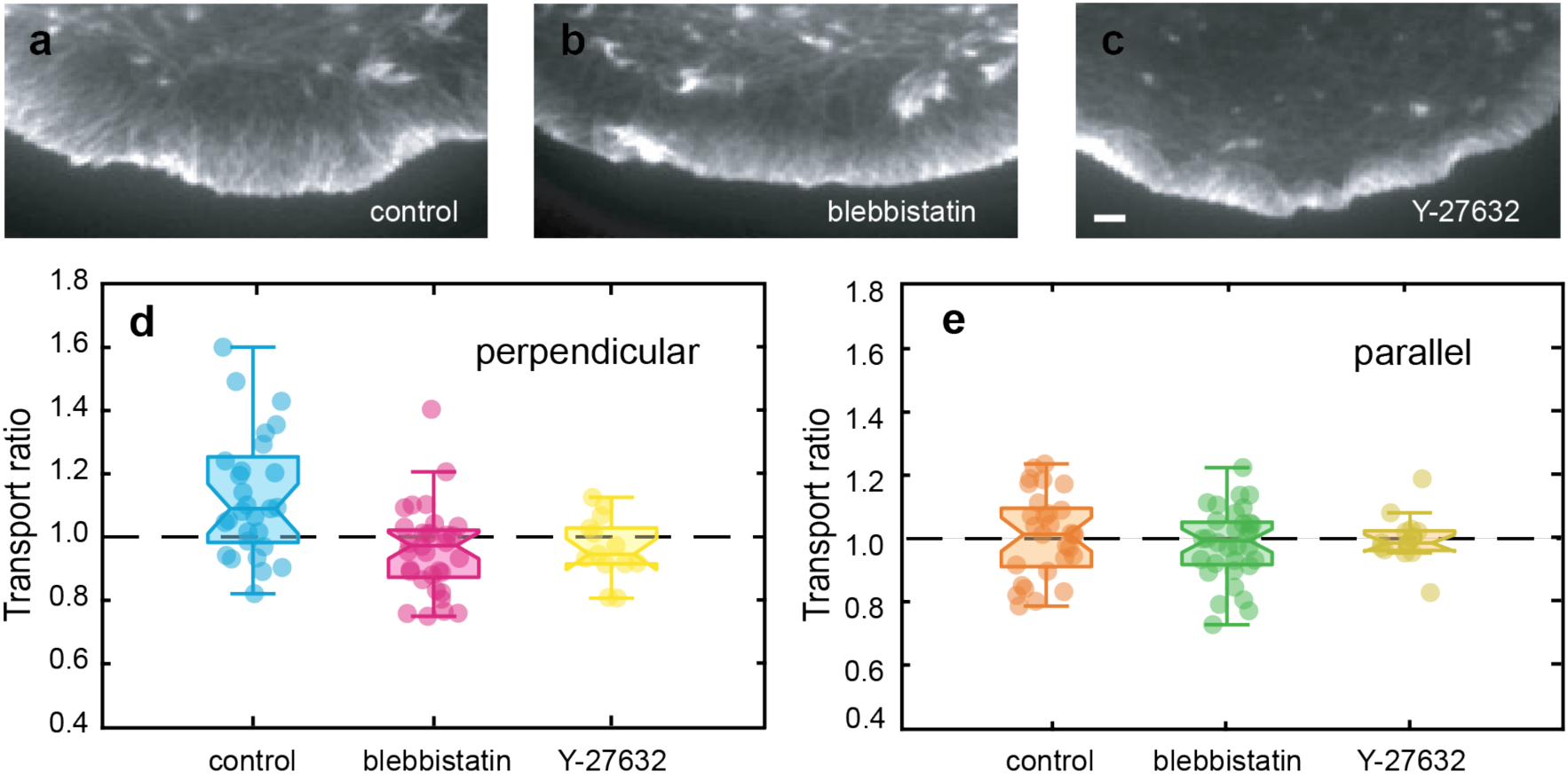
Comparison of Rho-kinase and myosin II ATPase inhibition. **a-c,** TIRF images of control CAD cells, cells treated with nitro-blebbistatin (0.2 μM), or Y27632 (1 μM), for 10 min before fixation and labeling with Alexa 647 phalloidin. The control cell has a wide lamellipodial branched network, followed by parallel fibers and transverse actin arcs at the border with the cell body. Nitro-blebbistatin-treated cells have a shorter lamellipodium, fewer parallel fibers, and a shallower arc. Y27632 treated cells have a shorter lamellipodium, followed immediately by transverse actin arcs. **d-e,** Transport ratio for control and blebbistatin treatments from Fig 2e, reproduced here for comparison with Y27632, n=13. P values from Scheffe post-ANOVA show no significant difference between blebbistatin and Y27632 treatments either parallel or perpendicular to the edge and no significant difference between the conditions parallel to the edge. Scale bar, 200 nm.

**Extended Data Fig 3.**
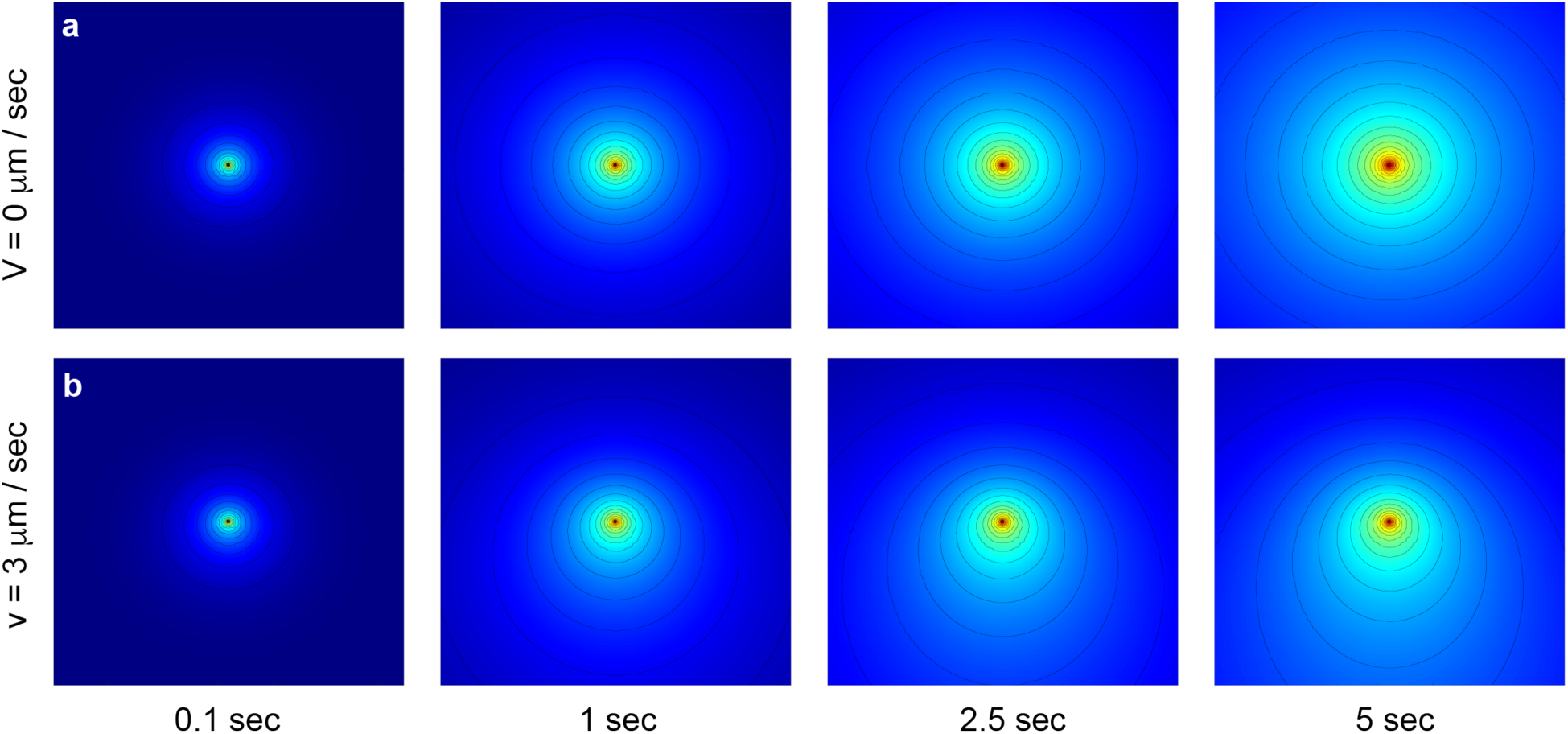
Mathematical simulation of distribution from a point source with and without fluid flow. **a,** Visualization of the analytical solution for continuous injection of material dissipating from a point source with a diffusion coefficient of 12 μm^2^/s, eqn (1) in methods. **b**, Simulations repeated with a downward directed fluid flow of 3 μm/s. The asymmetry is apparent after 1s. All images are 5 μm X 5 μm.

**Extended Data Fig 4.**
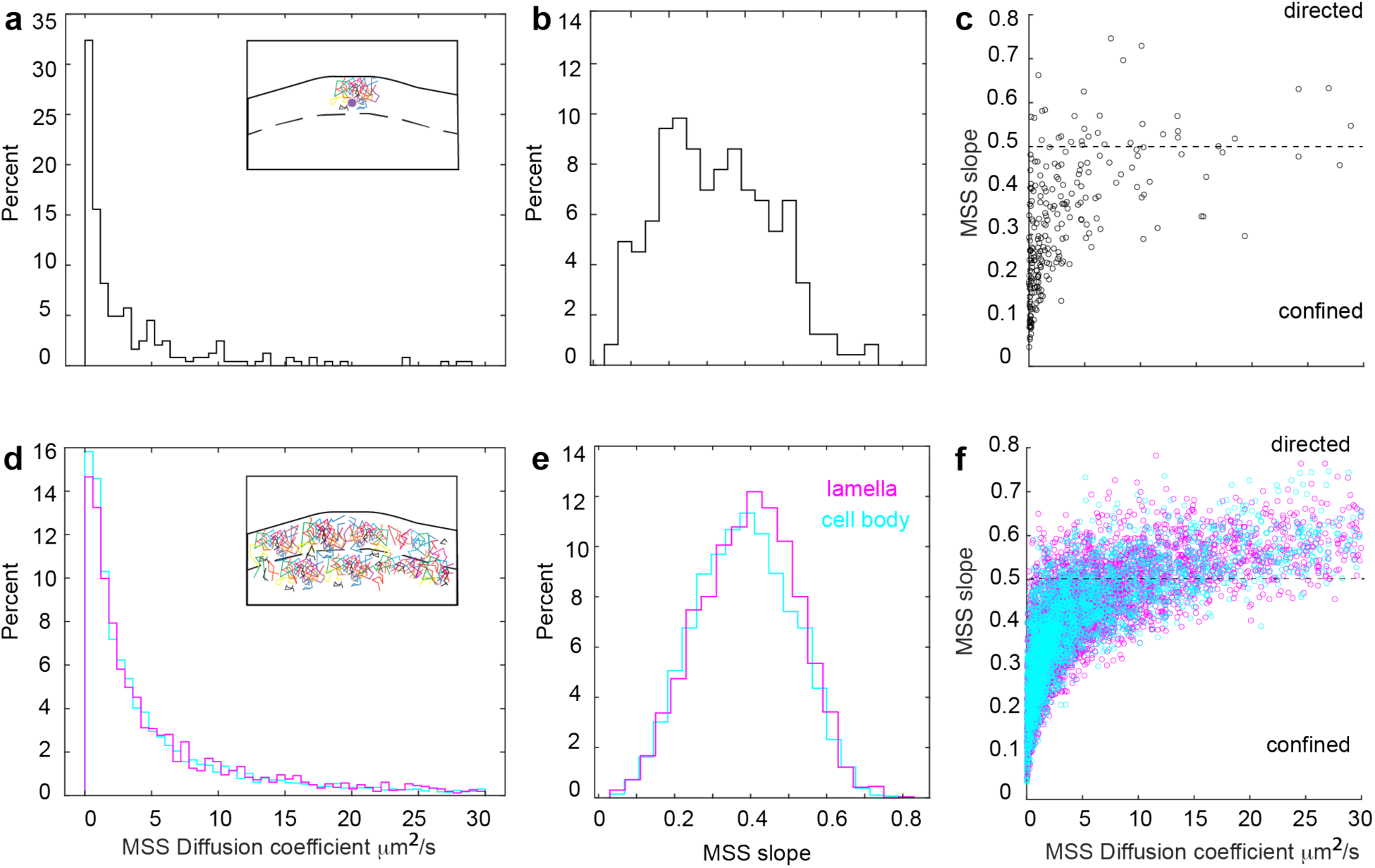
MSS analysis of single-molecule trajectories. **a-c,** Spot activation of mEos3.2 R62D in a single cell, 1451 trajectories analyzed. **d-f,** Halo G13R JF549 from 10 cells, 7831 trajectories analyzed, 2634 lamella, 4747 cell body**. a, d,** Diffusion coefficient of individual trajectories computed using the second moment (MSD). **b, e,** Slope of the Moment Scaling Spectrum (MSS) for the first seven moments. The slope is indicative of the mobility mode. A value of 0.5 defines random Brownian movement. Values above 0.5 indicate directed motion, and values below 0.5 indicate confinement ^29^. **c, f,** MSS slope as a function of diffusion coefficient.

**Extended Data Fig 5.**
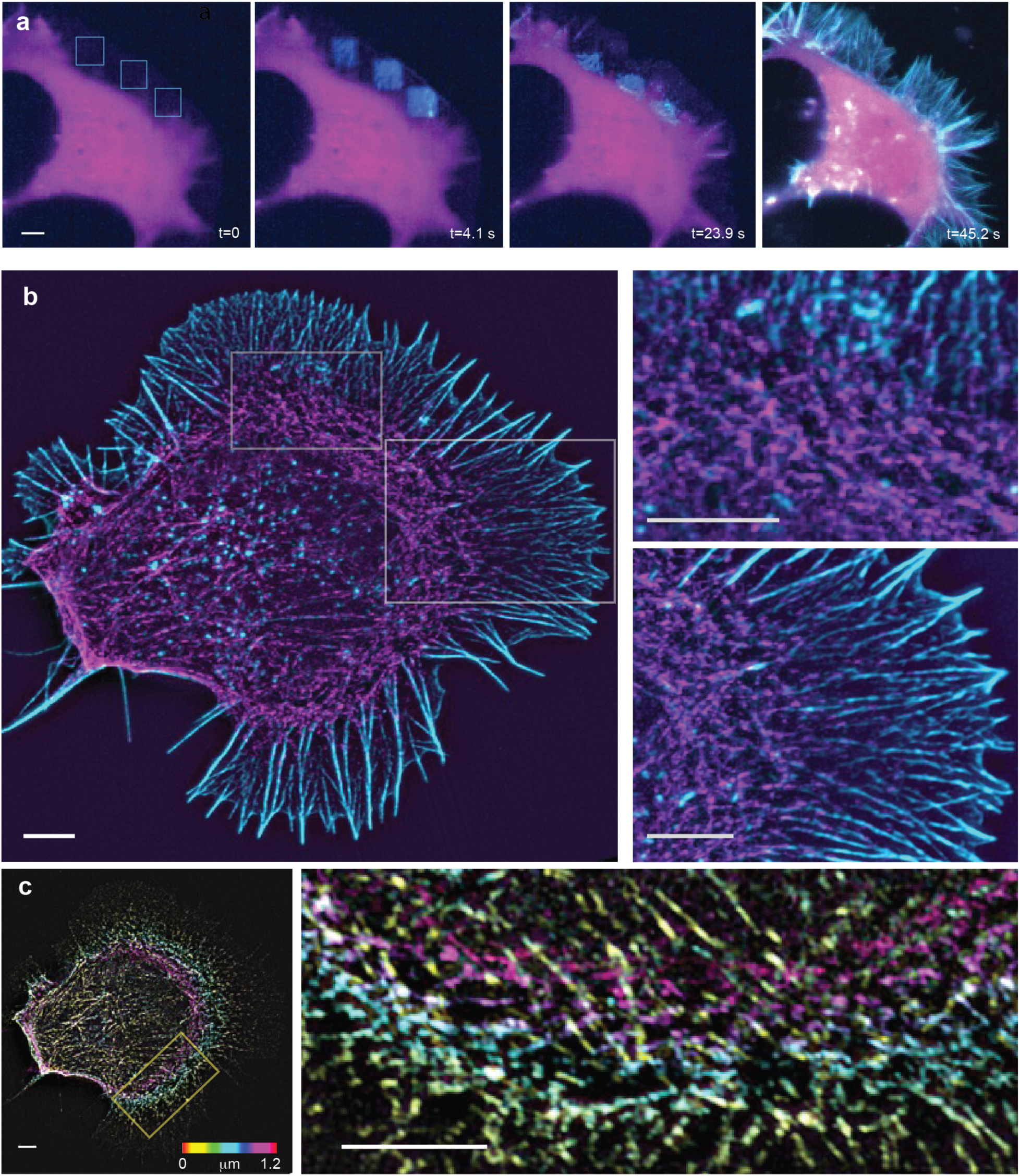
The myosin wall also disassembles the actin network in cells with filopodial-based growth cones. **a**, NG108 cells were transfected with PaGFP Actin and untagged RFP. Actin is not visible before photoactivation, t=0s. Immediately after photoactivating three square ROIs t=4.1s, actin is visible in each ROI, and a thin band of repolymerized actin is already visible at the leading edge. At 23.9 s, the ROIs have moved rearward due to retrograde network flow, and the bottom corners of the squares have disappeared as the network is depolymerized. The entire lamellipodium is also faintly illuminated. In the last panel, the entire cell was reactivated to view the entire network. **b,** 3D maximum projection SIM images of NG108 cells transfected with Emerald MLC (magenta), fixed and labeled with Alex 647 phalloidin (cyan). As with the CAD cells (Fig 3c), myosin is largely excluded from the lamella and concentrated at the rear of the lamella border. **c,** The same z-stack is color-coded for height, illustrating the vertical height of the actin-myosin wall. All SIM images represent at least three independently performed experiments. Scale bar, 10 μm.

**Extended Data Fig 6.**
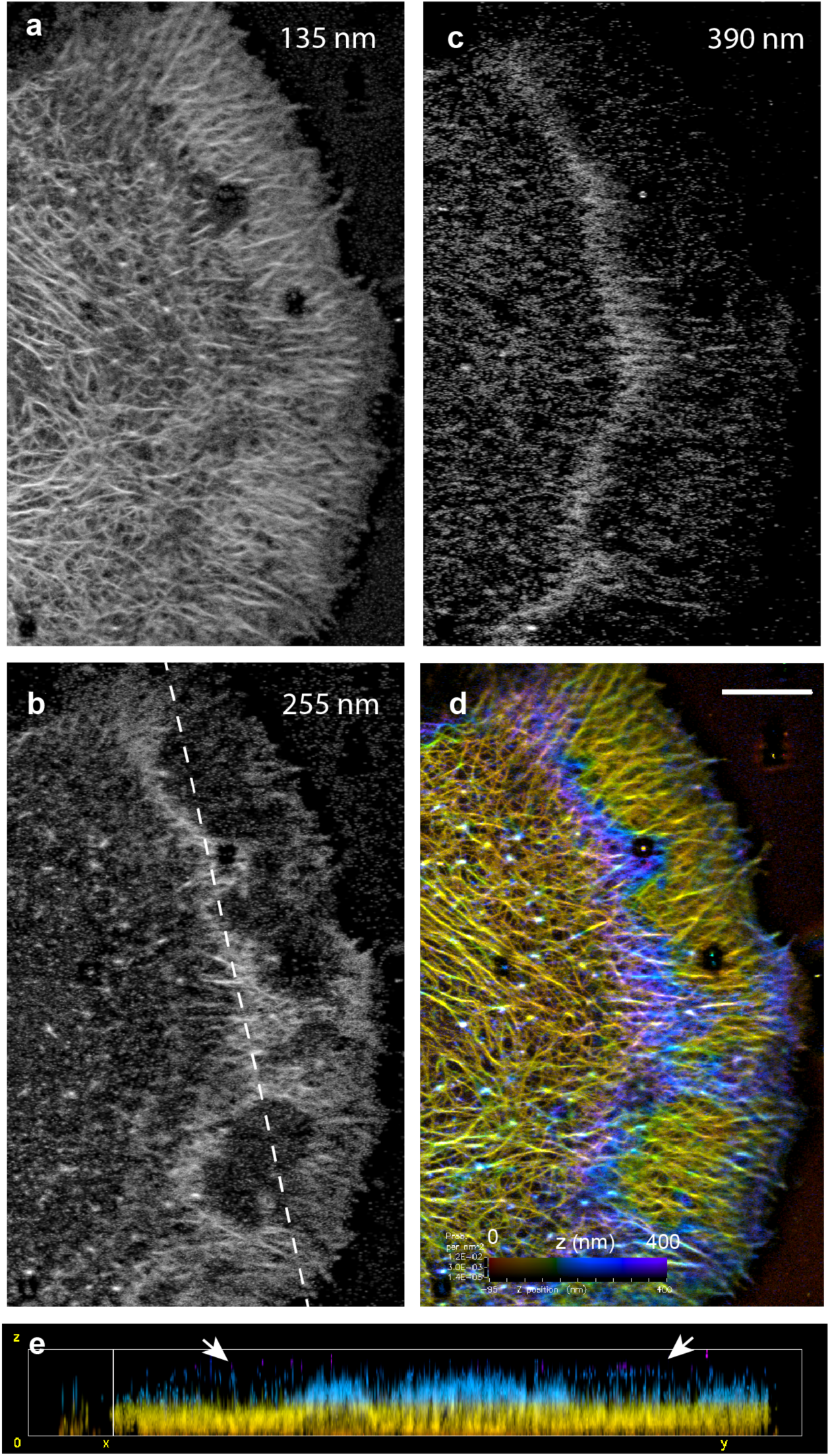
iPALM reveals that the wall creates a separate compartment in cells with filopodia-based growth cones. **a,** Virtual 15 nm thick optical x-y section of an NG108 cell imaged with iPALM and rendered with PeakSelector ^36^. The pseudo-color for the section matches the axial color bar in (**d**). **b,** a higher axial optical section illustrating compartments (dark holes) throughout the lamella. **c,** a higher plane showing a thin covering above the compartment. **d**, color-coded maximum intensity projection. **e**, y-z section taken through dashed line in (**b**) with the arrows pointing to the plane shown in (**c**).

**Extended Data Fig 7.**
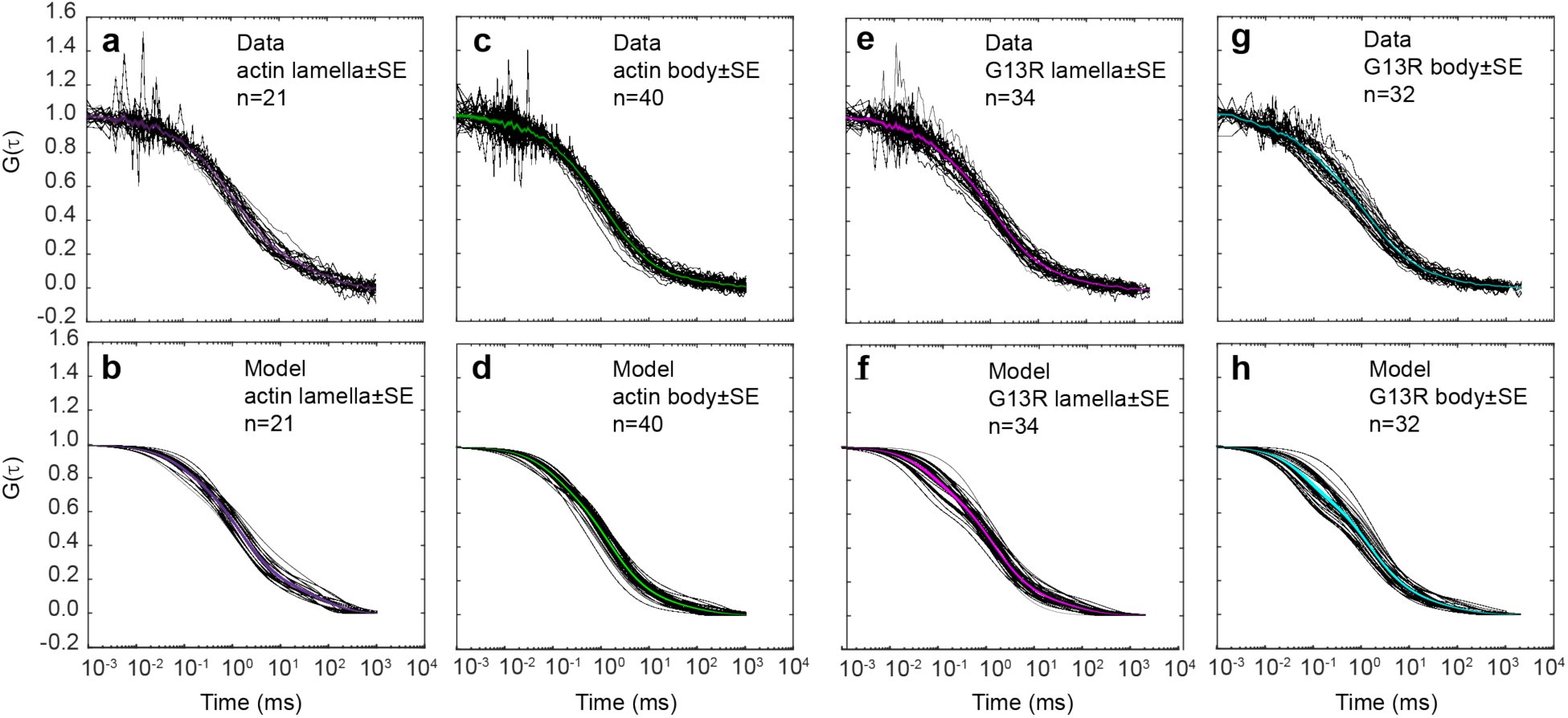
Transport of actin and non-polymerizable mutant measured in lamella and cell body by FCS. **a,** Normalized autocorrelation functions for Halo-tagged actin taken in the lamella. Each trace, as well as the mean ± SE, is depicted. **b,** Mean ± SE for model fit described in supplemental methods. c, d, Halo-actin in the cell body. **e, f,** Halo-G13R in the lamella. **g,h,** Halo-G13R in the cell body. All experiments were performed in transiently transfected CAD cells and labeled with JF549. Data represent at least three independently performed experiments. Calibrations were performed as in ^30^.

**Extended Data Fig 8.**
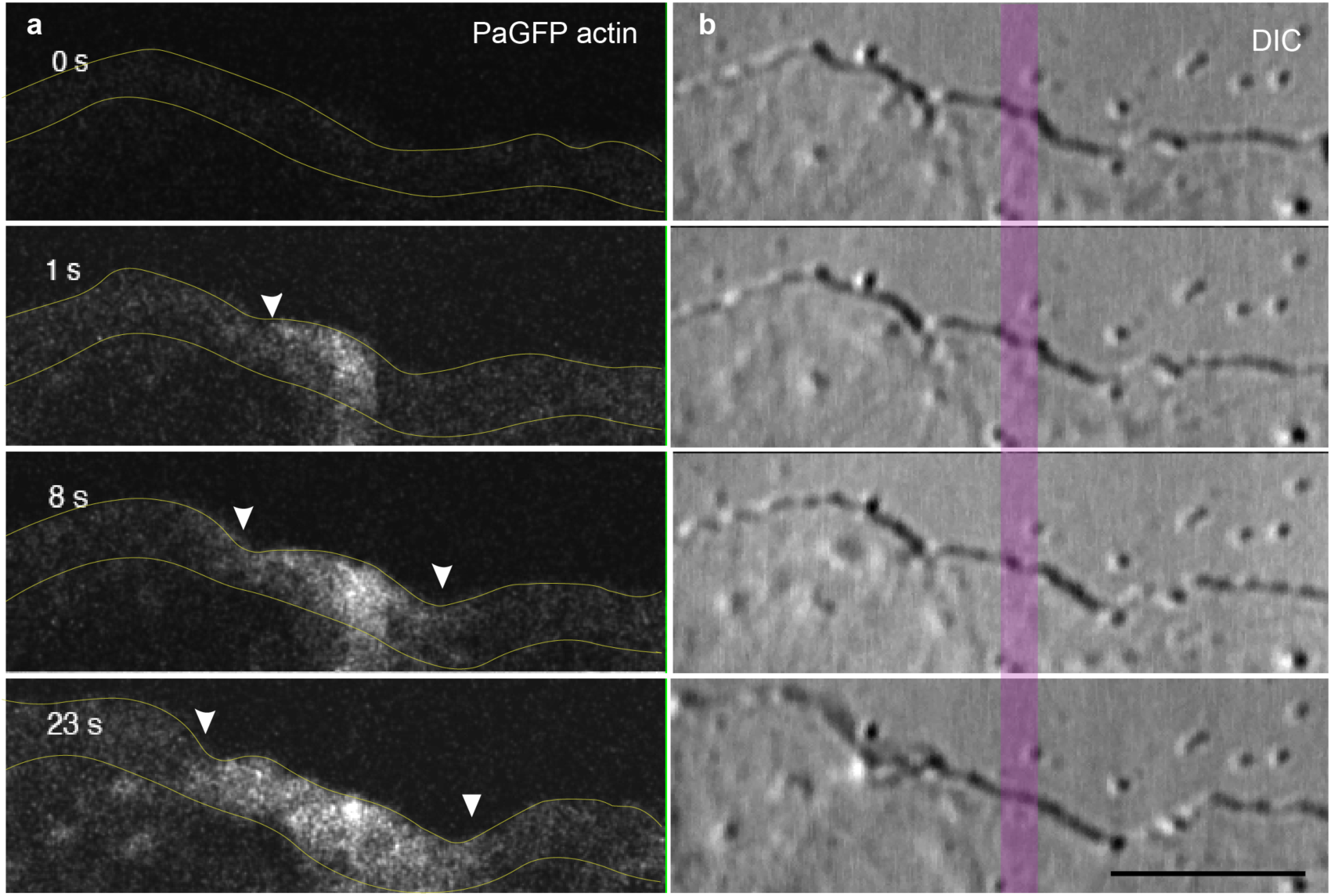
Compartmentalized flow in lamella switched direction as different sides of the cell advance. **a,** NIH 3T3 cell expressing PaGFPactin was photoactivated in a 1 μm wide stripe parallel to the leading edge between 0 and 1 s (magenta line in (**b**)). The fluorescence initially moves to the left of the stripe (3 s, arrowhead), where the DIC images in (**b**) indicate the cell is advancing. At 8 s, the cell advances slightly on the left of the stripe, and fluorescence appears as this edge moves forward (arrowhead). Fluorescence intensity indicates that the majority of actin remains in the lamellipodial region. Lamellipodium is indicated by thin yellow lines in (**a**). Scale bar, 5 μm.

## Methods

### Coverslip Cleaning and preparation

For single-molecule, structured illumination, fluorescence correlation microscopy (FCS), and photoactivation experiments, number 1.5 coverslips were purchased from Warner Scientific (25 mm) and sonicated for 30 min in Luminox (Alconox). Following three washes in approximately 300 ml of Milli-Q water, the coverslips were rinsed in acetone, air dried, and then backed dry in a 110°C oven for 45 min. Coverslips were cooled to room temperature, and then the plasma was cleaned with a Harrick plasma sterilizer. Coverslips were baked again at 110°C for 45 min in a covered crystallizing dish. 0.75 ml of HMDS (Sigma), which was stored under argon gas, was introduced into the center of the dish to vaporize and silanize the coverslips.

After an additional hour of baking, coverslips were cooled and stored in airtight containers until use. Coverslips were mounted in AttoFluor Chambers (Invitrogen) or CAMPO Chambers (LiveCell Instruments). MatTek dishes (MatTek Life Sciences) were used for duo scan experiments. For iPALM experiments, the number 1.5 coverslips were processed to embed gold nanorods to act as fiducials for drift correction according to previously published protocols ^36^. These coverslips were then cleaned and silanized as described above. All coverslips or dishes were coated with 40 μg/ml mouse laminin (MP Research) or 5 μg/ml human plasma fibronectin (MP Research) at 4°C overnight before use.

### Microscopy

All live cell microscopy was performed at 37°C using a WPI Air Therm PID temperature controller, except for the FCS experiments, which used a Tokai Hit stage top incubator with 5%CO_2_. All experiments were performed in phenol red-free growth media specific for the cell line, supplemented with 15 mM HEPES.

#### TIRF and SMLM

A TIRF microscope was constructed around an Olympus IX-71 stand using a 60X1.49 NA objective with Coherent 405, 488, 561, and 637 nm lasers used to build in an open-air laser sled ^37^. An ANDOR iXion 897 EMCCD camera, under the control of Solaris software, was used for image acquisition. The TIRF angle was controlled by a programmable Thor Labs motor (Z812B) and custom LabView software. The AOTF (AA Optoelectronics) and the lasers were controlled with AA MDS and Obis software, respectively. The filter set was Semrock Penta set, Di01-FF409/49/57/652/759, 432/514/595/681/809.

#### Multi-color TIRF

A custom-built electronic circuit accomplished the frame switching between 561 and 637 lasers on the TIRF microscope ^37^. Results were denoised using the Fiji CSBDeep Noise2Void plugin ^38^, with training consisting of 30 epochs, patch size = 64 and a neighborhood radius of 5.

#### Spot-activation TIRF

A second Coherent 405 laser, equipped with a fiber optic, was introduced into the fluorescence light path using a Semrock FF458-Di02 dichroic mirror, creating a diffraction-limited spot in the image field of the standard TIRF microscope.

#### Structured Illumination Microscopy (SIM)

Structured illumination experiments were performed using a Zeiss Elyra 7 with 488 and 642 nm lasers and a Plan-Apo 63x/ 1.4 NA Oil DIC objective. Filterset#1 BP 57-=620 + LP 655, Filterset#2 BP 420-480 + BP 495-550.

#### Zeiss Duo Scan

NG108 stripe bleaching and box activation experiments. Images for bleaching experiments were collected at 10 frames per second (Fig 1) and 2 frames per second for box activation (Extended Fig 5). Images collected with the Zeiss Duo Scan were filtered with a Gaussian blur (σ=0.6) followed by an unsharp mask (σ=0.75). In addition, the photobleaching of time-lapse movies was corrected using the exponential fit option in Fiji ^39, 40^.

#### iPALM

The iPALM microscope and its usage have been previously described ^36^. Single-molecule images were rendered in Peak Selector using Group Peaks ^36, 41, 42^. X-z and y-z views were created using the Volume Viewer plug-in in Fiji with a 10X scale in z ^40^.

#### Fluorescence Correlation Spectroscopy (FCS)

FCS curves were collected using a Leica TCS SP8 Falcon FLIM/FCS microscope controlled using Leica LAS X (3.5.7.23225) and LAS X FLIM/FCS (3.5.6) software. Imaging was performed with Leica APO 86X / 1.20 NA water-immersion objective with a motorized correction collar (mottCorr, Leica). Prior to FCS measurements, the mottCorr was adjusted using *x-z*-scans in reflection mode to yield the sharpest cellular image.

### Cell Culture and Transfection

#### Cell lines and culture

CAD (Cat.-a-differentiated) cells (Sigma-Millipore) were grown in DMEM/F12 media containing 2 mM glutamine (and supplemented with 8% fetal calf serum NG018-15 cells (ATCC) were grown in DMEM containing 1X HAT supplement and 10% fetal calf serum. NIH 3T3 cells were grown in DMEM High glucose supplemented with 10% fetal bovine serum. All media and supplements were obtained from Gibco; serum was obtained from Cytivia-Hyclone.

#### Labeling

Cells were labeled for FCS by incubating with JF549 ^43^ or JFx650 ^44^ at a concentration of 200nM for 30 min. They were then rinsed twice in complete media before imaging.

#### Fixation

All cells were fixed at 37°C with 2% paraformaldehyde in PHEM (pH 6.8) ^45^. After five minutes, the fixation media was replaced, and the fixation process continued for another 35 minutes. Cells were rinsed 3X for 5 min each in PHEM before labeling with Alexa 647plus phalloidin (Invitrogen).

#### Transfection

NG108, CAD, and NIH 3T3 cells were transfected using Invitrogen LTX Plus according to the manufacturer’s protocol. After 3 hours, the transfection media was exchanged for complete growth media, and the cells were imaged the following day.

#### Constructs

EGFP-Actin (Clontech cat #6116-1). Apple, mEos3.2, and Emerald actin were gifts from Michael Davidson. (Addgene nos. 54862, 57446, and 53978, respectively). Halo-actin was a gift from Kia Johnson.

PaGFP-Actin was constructed by excising the color from PaGFP-C1 (gift from George Patterson) and EGFP-Actin ^23, 34^.

mEmerald-LC-Myosin-N7 was a gift from Michael Davidson (Addgene no. 54146). mNeon LC-Myosin-N7 was constructed using the enzymatic digests described in the Supplementary Information.

mEos3.2 ARC3 was constructed from EGFP-ARC3, a gift from Matt Welch (Addgene no. 8462)^46^, and mEos3.2-N1 (Addgene no. 54525) using SmaI and Not I.

mEos 2 Paxillin-22 was a gift from Michael Davidson (Addgene no, 57409).

pENTR-NLS-actin-R62D (Addgene no. 11831) was used to construct the mEos3.2 actin-R62 plasmids using Sali-HF and BmbG1 to combine the mutated region of actin with existing mEos3.2 actin plasmids. All other colors were constructed using enzymatic digests described in Supplementary Information. The insert regions of the newly constructed plasmids were verified by sequencing.

YFP NLS Beta-Actin G13R (Addgene no. 60615) was used to construct the mEos3.2 actin-G13R plasmid using SalI-HF and BamH1 to combine the mutated region of actin with existing mEos3.2 actin plasmids. All other colors were constructed using enzymatic digests described in Supplementary Information. The insert regions of the newly constructed plasmids were verified by sequencing.

### Analysis

#### Transport Ratio

Baseline images were collected before initiating activation (typically 50-75 frames). These images were averaged, and the result was subtracted from each image frame in the experimental time series. After activation, the first 500 frames (1000 frames for Arp3 and paxillin) were summed to create an image that is the physical solution to eqn. (1), with t=7.5 s. This image was contrast stretched, and the intensity was recorded through a 5-pixel-wide line, 10 μm long drawn orthogonal to the leading edge and through the center of the activation spot or a 5-pixel-wide line drawn parallel to the edge and through the spot center.

Software written in Matlab normalized each of these intensity curves, smoothed them with a spline fit, detected the peak (center of the activation spot), and integrated the area under the curve on both sides of the center. These two area calculations were then compared. The ratio of these areas (leading edge/cell body or parallel side1/parallel side2) was defined as the transport ratio. Since there was no morphological rationale for the side-to-side calculation, the choice of which side was “leading” or “body” was randomized.

In the presence of fluid flow, the areas of the two halves of the curve drawn orthogonal to the leading edge should not be equal. If the area of the half of the curve closer to the leading edge is divided by the area closer to the cell body, then the ratio should be greater than 1. However, the fluid flow would not be expected to affect the lateral distribution of mass in the direction parallel to the leading edge. In this direction, the ratio is expected to be 1.

#### Transport Ratio Simulation

Plume profiles in Extended data Fig 3 were generated in Matlab by numerically integrating the solution for a continuous injection at the origin into a steady plane flow in the x-y plane: (equation 10.6.38^22^):

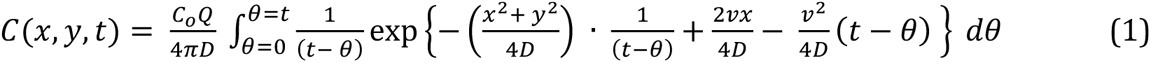

Where *C_o_Q* is a constant injection rate of material, *D* is the diffusion coefficient, and *v* is the velocity of the flow in the x direction.

#### Localization and Tracking

All single-molecule tracking was performed with TrackIt^47^

#### FCS

The FCS curves were exported to Matlab, where the autocorrelation curve was fitted using a two-component model that accounted for both diffusion and flow, with the flow affecting both species equally^31, 32^

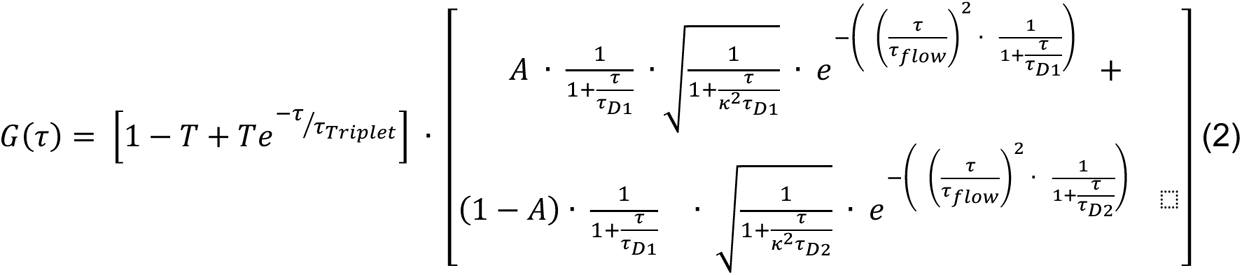

Where *T* is the triplet state fraction, τ_triplet_ is the triplet state lifetime in ms, τ_D1_, τ_D2_ is the diffusion times of species 1 and 2 in ms, *A* is the fraction of molecules with diffusion time τ_1_, τ_flow_ is the flow time in the volume in ms, and κ is the structure parameter of the focal volume (κ = 5). The diffusion coefficient was calculated by: 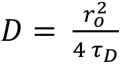 and the velocity by: 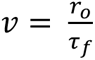 (*r*_o_ = 0.2543 μm). All traces were analyzed individually.

#### Peclet Number

The Peclet number is defined as the ratio of the advective transport rate/diffusion transport rate. For mas transport it is given as:

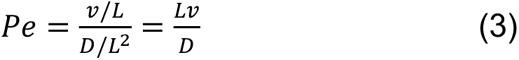

Where *v* is velocity, *L* is the characteristic length, and *D* is the diffusion coefficient. When the Pe number is greater than one, advection dominates, and when Pe is less than one, diffusion is the dominant form of transport.

